# Exploring DNA movement through the application of droplet based high efficient chromatin conformation capture (DropHiChew) and loop velocity

**DOI:** 10.1101/2024.06.26.600744

**Authors:** Chen Zhang, Yeming Xie, Chen Tan, Zhichao Chen, Mei Guo, Chong Tang

## Abstract

This study presents a novel approach to understanding DNA movement dynamics through the development of a droplet-based, high-efficient chromatin conformation capture method, known as DropHiChew, and a new algorithm, loop velocity. DropHiChew, a user-friendly and cost-effective technique, employs the 10X single-cell systems allowing for easy experimental implementation. The loop velocity algorithm, on the other hand, enables the estimation of the speed and direction of cell development, providing a dynamic perspective on chromatin movement. Even with shallow sequencing, our loop velocity algorithm accurately gauges the trajectory of DNA motion and cellular states. The combined use of DropHiChew and loop velocity offers potential for a wide array of applications in future chromatin capture studies, including disease modeling, cellular differentiation studies, and developmental biology.

## Introduction

Chromatin conformation refers to the way DNA is arranged in three dimensions within the nucleus of a cell. It plays a crucial role in controlling gene expression and other cellular processes. Understanding chromatin conformation is important because it helps us understand how the genome is organized and how different regions of DNA interact with each other. Studying chromatin conformation is significant in the context of both disease and biological research. Abnormalities in chromatin conformation have been linked to various diseases, including cancer and genetic disorders^1^. By unraveling the complexities of chromatin conformation, researchers can gain a better understanding of the underlying mechanisms driving these diseases and potentially discover new treatment targets.

Scientists have been diligently studying the dynamic folding of the genome within the cell nucleus over the years. It’s been found that DNA is extruded into loop structures with the help of CTCF and cohesin^2^. This ongoing loop extrusion forms a more complex structure referred to as a topological association domain (TAD)^3^. The genome is then organized based on its proximity to the nucleus membrane, forming the constitutive lamin-associated domain (cLAD) and constitutive inter-lamin-associated domain (ciLAD). Notably, these domains have shown a strong correlation with the AB compartment in HiC experiments^4^. However, many of these structures are defined from a static perspective and average phenotype bulk sample. Given that DNA is highly dynamic within cells, we aim to observe the genome from dynamic viewpoints, which will greatly broaden our understanding of genomes. To investigate the dynamic actions of the genome, we require single-cell resolution to estimate movement in each cellular state.

Addressing a key challenge, scientists have created a method named single nucleus HiC (scHiC). This groundbreaking technique enables the examination of chromatin conformation at the level of individual nuclei, which helps reconstruct the 3D structure of chromatin. The first version of scHiC featured biotin enrichment to amplify interactions, using a procedure similar to conventional HiC^5^. This involved genome digestion using a particular enzyme, adding biotin into the ends, and connecting proximal fragments. Paired-end sequencing was then applied to identify interactions between these fragments. It was later found, though, that biotin enrichment in single nuclei led to fragment loss, which impacted the resolution of observed chromatin conformations.

In response to this challenge, researchers developed a method known as Dip-C^6^. This approach eliminates the requirement for biotin incorporation and purification, enabling whole-genome sequencing without any DNA loss during the purification process. However, the solution does impact sequencing depth, resulting in over 90% of sequencing data being unutilized and only 8% of data providing valuable contact. In fact, most single-cell 3C techniques in common use today are based on the Dip-C concept.

Our laboratory recently developed snHiChew, a cutting-edge method using methyltransferase to label the ligation scar (GATC pattern) after PCR in Dip-C, effectively boosting the efficiency to 45% valid pairs - currently the highest efficiency in single-cell chromatin capture technology. While impressive, these techniques require labs to design their cellular barcodes and apply the split-pool method for cell labeling. Therefore, we’re on a quest for a more straightforward and cost-effective approach that can yield positive results for the majority of labs.

As we fine-tune our experimental procedures, it’s equally important to develop an algorithm that interprets DNA motion and sets parameters around velocity for a dynamic view of the genome. This will enhance our grasp of the biological impacts of chromatin movement, such as shifts in the AB compartment and TAD transitions. More crucially, understanding genome velocity could lead to more accurate forecasts of DNA’s future position, which can help determine the upcoming cellular state. In contrast, RNA dynamics have a well-documented history ^7^. Both our team and others in the field typically use intron-rich RNA to signify RNA synthesis, and exon-depleted RNA to denote RNA degradation. This method allows us to gauge the rate of individual RNA production and degradation, which could be used to predict cellular fate.

Although using introns as indicators of nascent RNA speed is indirect and potentially biased^8^, it’s a necessary approximation at this point. However, the scientific community is continuously seeking to refine algorithms and methodologies for a more accurate estimation of nascent RNA expression^9–14^.

On the other hand, measuring DNA movement gives us a direct insight into the distance travelled from one location to another. By using a differential equation over time, we can get a more precise speed estimate. This could potentially be the first accurate distance-based velocity estimation in single cell studies, if confirmed. So far, there’s no existing research on how to calculate DNA movement velocity.

Our research is focused on finding practical and cost-effective ways to extend our knowledge of the DNA dynamic movement. We’ve put to use the 10X single-cell systems to develop a user-friendly, droplet-based chromatin conformation capture technique called DropHiChew. This efficient process involves digesting single nuclei with enzymes, ligating nearby fragments, attaching adaptor sequences using Tn5, and packaging single cells with barcode gel beads via the 10X chromium system. After this, we carry out in-droplet reverse crosslinking and label each nucleus DNA with a barcode through linear amplification. Given that the 10X single cell system usually yields thousands of cells, a budget of 2M per cell could be beyond the budget for many labs. To capture the movement of DNA in depth, we’ve crafted a new algorithm, loop velocity, based on loop extrusion models. Our research indicates that even with shallow sequencing (50k contacts per cells), we can accurately gauge the speed and direction of cell development using our loop velocity algorithm. We’re confident that this method has great potential for diverse applications in future chromatin capture studies.

## Results

### Experimental design of DropHiChew

Droplet-based platforms like 10X Genomics and MGI C4 are leading the way in the single-cell field, bringing these experiments within reach for many labs. However, these platforms haven’t been used for single cell chromatin conformation capture (3C) experiments yet, which makes single cell 3C less accessible for most researchers. Considering the cost of sequencing the whole genome in Dip-C, we aim to enrich the valid pairs after PCR. This way, we can maintain a balance between capture sensitivity and cost-effectiveness. To achieve this, we plan on using dam methyltransferase to label ligation scars (GATC pattern, digested by DpnII in previous step), then enrich the ligation scar by methylation immunoprecipitation (cite). But, there is a significant constraint when using this approach on commercial systems. The adaptor sequence or barcode sequence needs to avoid the GATC pattern, otherwise, these sequences will also be captured during the enrichment step, leading to a considerable barcode bias. Given that 10X Genomics uses random sequencing, 5% of barcodes have GATC, and MGI C4 has around 7% of barcodes containing GATC. This high prevalence of GATC-containing barcodes poses a challenge for implementing HiChew on commercial platforms.

We’ve developed a solution to this problem with our DropHiChew design(Figure1.A). After in situ genome digestion by DpnII (GATC pattern) and promaxite ligation (ligation scar-GATC pattern), the cells are transposoned using the Tn5 adaptor in the kits (step 1-3). These cells are then loaded into the 10X platform with the scATAC-seq program (step 4).

GEL beads, which carry barcoded primers, encapsulate the cell within a droplet(Figure1.A). A linear PCR is then performed to attach the barcode primer to the adapted DNA fragments. After the droplet has been emulsified, we amplify the DNA library using PCR to guarantee a sufficient amount of DNA.

We use a unique technique to increase the concentration of the 3C ligation scar (GATC) (Figure1.A). This involves employing two universal primers in PCR, one with several phosphorothioate bonds and another without. These bonds are resistant to exonuclease digestion. Subsequently, we use the T7 exonuclease to break down the plus strand DNA, leaving the minus strand DNAs intact. The another primer that avoids the barcoding region is then used to extend on the single strand DNA and create a double strand DNA with a single strand barcode at the end (partial extension) (step 5-6). The dam methyltransferase (GAmTC) can only act on the double strand DNA, allowing the single strand barcode to escape the methyltransferase modification (step 7). As a result, only the ligation scar with GATC can be modified. We then carry out methylation-specific immunoprecipitation to enrich the GATC ligation scar. Finally, we assess the DropHiChew design to confirm if it successfully enriches the fragment with the ligation scar.

### Evaluation of the DropHiChew

In order to assess the effectiveness of DropHiChew on single cell differentiation, we conducted a mix of HEK293 cells and mouse 3T3 cells in equal proportions(Figure1.B). Upon final analysis, we identified 1129 cells as human and 1610 cells as mouse, with 113 cells classified as doublets. This doublet rate aligns with the 10X parameter of a 0.04 collision rate.

We’ve performed further deep sequencing of individual cells in HEK293, improving our validation process. This resulted in a remarkable 6864 cells per reaction, 100K valid contacts per cell, and 24K unique valid pairs (dedup) per cell, with a duplication rate of 74% (Sfig1.A). The cis-trans ratio was 1. After debarcoding, we examined the potential for a GATC bias in barcode detection due to GATC enrichment. But our design is ready for this. Our investigation revealed the GATC barcode accounted for 5%, which is consistent with the percentage of the 10xATAC published whitelist (Figure.1C). This implies that the capture bias in DropHiChew is not considerable. For a deeper understanding, we compared GATC barcodes with control AGCT and CATG barcodes. GATC barcodes only had a 21.4% higher read number than control AGCT, CATG barcodes, which is a considerable advancement from our control method that was fully methylation labeled without avoiding barcode (78%) (Sfig.1B). In addition, when comparing the ratio of valid pairs, which indicates the enrichment efficiency on these different barcodes, the difference was minimal, suggesting similar enrichment efficiency with or without GATC in barcode (Sfig.1B). All these observations indicate that DropHiChew doesn’t introduce a significant enrichment bias with GATC or non-GATC barcodes.

**Figure. 1.**
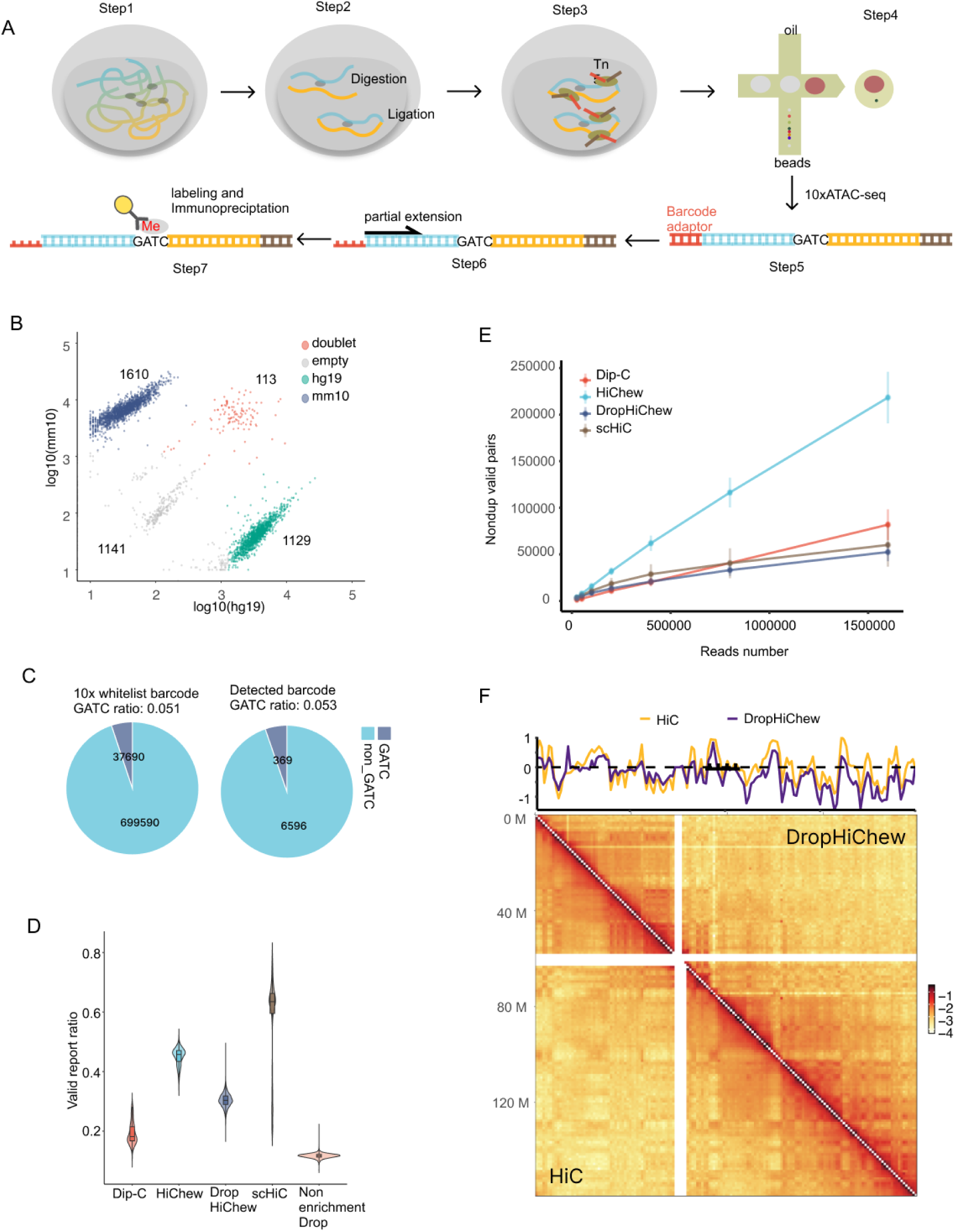
(A) We’ve outlined the DropHiChew experimental procedure in an illustration. (B) We conducted a mixture of HEK293 and CHO cells on the DropHiChew. The X and Y axis show the reads belonging to the mouse and human genome in each cell. (C) We’ve provided a pie chart that shows the percentage of the GATC barcode contained in the 10x whitelist, and the percentage of reads with GATC motif barcodes. (D) The violin plot displays the valid report ratio of different technologies. You’ll note that the non-enriched DropHiChew refers to the DropHiChew process without motif enrichment steps. (E) There’s also a line plot showing the number of unique valid pairs (deduplication) across different cells with varying sequencing depths. (F) We’ve included a contact heatmap on Chr7, comparing the DropHiChew and HiC.

When evaluating data usage efficiency of a 3C experiment, it’s standard practice to consider the valid pair ratio (valid pairs/qualified reads). This is a key measure of the experiment’s success. Here, snHiC delivers an impressive mean valid pair ratio of 63.4% (Figure1.D). Following closely, snHiChew presents a respectable 45.7% ratio. DropHiChew also performs well with a 30.3% ratio, while Dip-C shows a 18% valid pair ratio. In contrast, DropHiChew without ligation scar enrichment, which mirrors Dip-C performance on the 10X platform, yields only an 11.7% ratio. This could imply that the 10X droplet platform may affect chromatin capture efficiency compared to the plate-based Dip-C. It’s also worth noting that the unenrichment method can result in as much as 88% data usage wastage.

While our valid pair ratio is high, we must consider potential duplicate reads that aren’t useful for our analysis. After removing these duplicates, we mapped the unique valid pairs across different sequencing depths of individual cells (Figure 1.E). Both DropHiChew and snHiC generated up to 5 times fewer contacts per cell compared to HiChew at 1.6M sequencing reads. The performance of DropHiChew aligns closely with snHiC, reaching the plateau of unique valid pairs rapidly around 50K. This suggests that, like snHiC, DropHiChew might be missing a significant number of contacts, which could affect capture sensitivity. The majority of steps between HiChew and DropHiChew are the same, except for DropHiChew using Tn5 for adaptor attachment. Comparing the impact of Tn5 on crosslinked and non-crosslinked cells indicates that Tn5 might have difficulties efficiently attacking the crosslinked genome, potentially causing some DNA loss during the process (Sfig1.C).

We’ve taken a detailed look at DropHiChew’s performance by aggregating cell data as a simulated bulk sample and comparing it with HiC. The distance decay curves for both DropHiChew and HiC were notably similar (Sfig1.D). When we examined various resolutions on the contact heat map, from AB compartments to TAD, DropHiChew exhibited a correlation of 0.82 and 0.77 in compartment eigen value and TAD insulation score distribution respectively (Figure1.F, Sfig1.EF). Overall, DropHiChew’s results align well with the standards set by regular HiC experiments.

To sum it up, DropHiChew offers a viable and commercially sustainable solution for handling large quantities of cells, although it does have a restricted range of valid contacts per cell. Despite this, we’re confident that the limited contact depth won’t prevent us from tracking DNA movements. Consequently, we’ve developed a velocity algorithm that enables us to comprehend the dynamic movements of DNA, even with shallow sequencing.

### The loop velocity development and the TAD dynamic model

Recent studies ^15^ ^16,17^ have shown that changes in TAD insulation or other structures during gene expression can offer valuable insights into chromatin motion during cellular development. However, even if the DNA does move, the speed cannot be simply determined by dividing the distance by time due to unknown time factors. Thus, the velocity of DNA movement can only be estimated using mathematical models. Despite these advancements, our understanding of kinetic modeling of DNA still has room for growth.

The Hi-C matrix allows us to visualize structures like TADs as beads in a 3D model, showcasing their high interaction frequencies. TADs are known for these frequencies, particularly their distinct boundaries created by significant interaction overlap in cells within these regions. These TAD boundaries often coincide with CTCF binding sites, sparking a lot of academic interest^18^. This correlation, backed by experimental findings^2,19^, led to the development of the loop extrusion model.

In the loop extrusion model, SMC protein complexes like cohesin bind to DNA and start extrusion (Figure2.A). Here, DNA moves through these cohesin rings, forming a loop. This extrusion is limited on one side by the DNA’s initiation boundary, blocked by CTCF binding. The initiation boundary (I boundary) remains fixed, while the other stabilizing boundary (S boundary) moves towards the cohesin, until a loop structure forms. After this, the cohesin unloads, and the TAD disassembles. For example, in a selected region of 43MB-44MB and the Human Embryonic Kidney (HEK) cells, we observed interactions starting at the initiation boundary and gradually shifting to the stabilizing boundary (Sfig.2A). These observations further support the loop extrusion model.

Based on our observations and existing theories, we propose a hypothesis that variability suggests that the process of a TAD forming with loop extrusion theory within a cell can be likened to a domain generating, stabilizing, and disassembling within a set range (Figure1.A).

The hypothesis we’re examining provides a foundation for developing loop velocity. The subsequent inquiry is how to create the mathematical model. Previous studies have suggested that RNA velocity is estimated by a three-state model that includes RNA transcription (nascent unspliced RNAs), a steady state, and RNA degradation (degrading spliced RNAs)^7^. These three states of RNA velocity constitute what we refer to as the spindle model, with the steady state playing a vital role in this arrangement. Interestingly, these three RNA states align with the domain generation, stabilization, and disassembly states within TADs that we’ve discussed.

We’ve analyzed a sample region (Chr3:4203750∼43062500), and looked into the interaction distribution with both the initiation and stabilizing boundaries within each cell (Figure2.B). The data suggests that these cellular processes contribute to the formation of a spindle model. The upper red curve represents an increase in contact with initiation boundary over time, indicative of domain generation. Conversely, the lower black curve shows a progressive decrease in contact with the stabilizing boundary, indicating disassembly. The intersection point represents the stabilization phase, analogous to the steady state seen in the RNA model. This flawless spindle model paves the way for the development of a loop velocity algorithm.

We propose that the frequency of contact formation with the initiation boundary per unit time during loop extrusion is a constant, α (Figure2.A). As the loop continues to wind and roll away from the initiation boundary moving towards the stabilizing boundary on the other side, there is a decrease in contacts with the initiation boundary, a speed we’ve set to β.

At the initiation boundary, the decrease in contacts corresponds to a potential increase in contacts at the stabilizing boundary. Once the other stabilizing boundary with CTCF meets the cohesin, a stabilized TAD is formed. After this, the cohesin is unloaded from the DNA, and the DNA continues to unwind into space - the internal contacts decrease at speed γ.

From this one-sided loop extrusion process, we can derive the following ordinary differential equations (ODE):

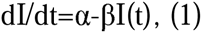

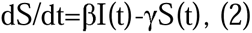

Here, α denotes the constant that generates contact at the initiation boundary, while β signifies the rate at which active contact is reduced at the initiation boundary during loop formation. Meanwhile, γ represents the rate of passive contact reduction. The term dI/dt refers to the interaction with the initiation boundary per unit of time, while dS/dt denotes the stabilizing boundary per unit of time. I(t) indicates the number of contacts with the initiation boundary at time point t, and S(t) corresponds to the number of contacts with the stabilizing boundary at the same time point.

To determine the unknown parameters α, β, and γ, we infer them once a stable TAD is established in a steady state. During this stability, the DNA loop extrusion and winding process halts, and the extrusion speed relative to the two edges approaches zero (Figure2.B). At this point, both dI/dt and dS/dt are zero. This parameter estimation process is akin to the one used in RNA velocity. We set β to 1, meaning we measure all rates in terms of the reduction rate. Following some equation transformation, γ turns out to be I(t)/S(t), indicative of the dot plot’s slope, which we can estimate via linear regression. We then solve for α. Having calculated all the parameters, we can estimate the subsequent cellular state through velocity calculation.

To sum up, you can find the derivation of the complete equation and parameters in the supplementary notes of RNA velocity and Region velocity, which make use of the ODE equation and the steady state resolution method. This velocity model, which we’ve decided to call Loop velocity, uniquely uses Hi-C TAD features and integrates the loop extrusion concept.

### Perform the loop velocity on the HEK293 cells from DropHiChew

Initially, we apply loop velocity to HEK293 cells from DropHiChew, examining the functionality of alpha and gamma factors for velocity prediction. We’ve identified xxx TADs from these cells, aiming to determine a relationship between loop movement and time. The cells’ time is represented by replication scores, which indicate progression from early to late replication stages. Following this, we compute average interactions within initiation and stabilization boundaries of selected TADs (Figure2.C, Sfig2.B). We observed a contrasting trend of both boundaries with replication states from early to late stages, indicating a significant correlation between interaction movement and development time. This strongly supports the spindle model previously discussed.

We then looked at the I_S dot plot for spindle models across different regions (Sfig2.C) and saw that 57% of TAD regions could form the model (Figure2.D), suggesting a potential link with time. This is a significant improvement over RNA velocity. The limitations of RNA velocity come from its dependence on short read sequencing from 10X scRNA, which hampers its ability to capture intron reads in RNAs. We’ve previously mentioned that this could lead to a considerable underestimation of introns and prevent several genes from forming the spindle-like model, which could distort RNA velocity predictions^8^. On the other hand, loop velocity aligns well with the spindle model and enables the analysis of more TADs (57%) (Figure2.E), enhancing prediction accuracy based on our observation.

We assessed the alpha gamma for each relevant TAD. Our model suggests that a higher alpha value points to a quicker establishment of contacts, while a heightened gamma value signifies a faster pace of TAD dissolution. Additionally, we delved into the possibility of alpha and gamma being associated with genome organization aspects.

Our research suggests that TADs in the A compartment typically exhibit a significantly higher alpha value compared to those in the B compartment (Figure2.F). A global correlation of 0.4 between alpha values and eigenvalues underscores this noteworthy relationship (Figure2.G). Furthermore, TADs with high chromatin accessibility and other markers of active histone also present significantly higher alpha values, aligning with the established correlation between eigenvalue and active transcription (Sfig2.D). This leads us to infer that alpha, indicative of TAD generation speed, may have a close association with eigenvalue and chromatin activity.

Interestingly, we observed that gamma, representing the speed of TAD dissolution, delivered contrasting results. It was considerably lower in the A compartment and active histone markers, albeit not as significantly as alpha’s correlation (Sfig2.D). The most robust correlation was seen with boundary strength, a factor that depicts TAD stability (Figure.2F). A correlation of −0.2 suggested that TADs with higher boundary strength dissolve more slowly, which makes sense. Further exploration of the boundary strength-related factor, CTCF, revealed a negative association with the gamma factor (Sfig2.D). This implies that the speed of TAD dissolution (gamma) has a weak negative relationship with both CTCF signal and boundary strength.

**Figure. 2.**
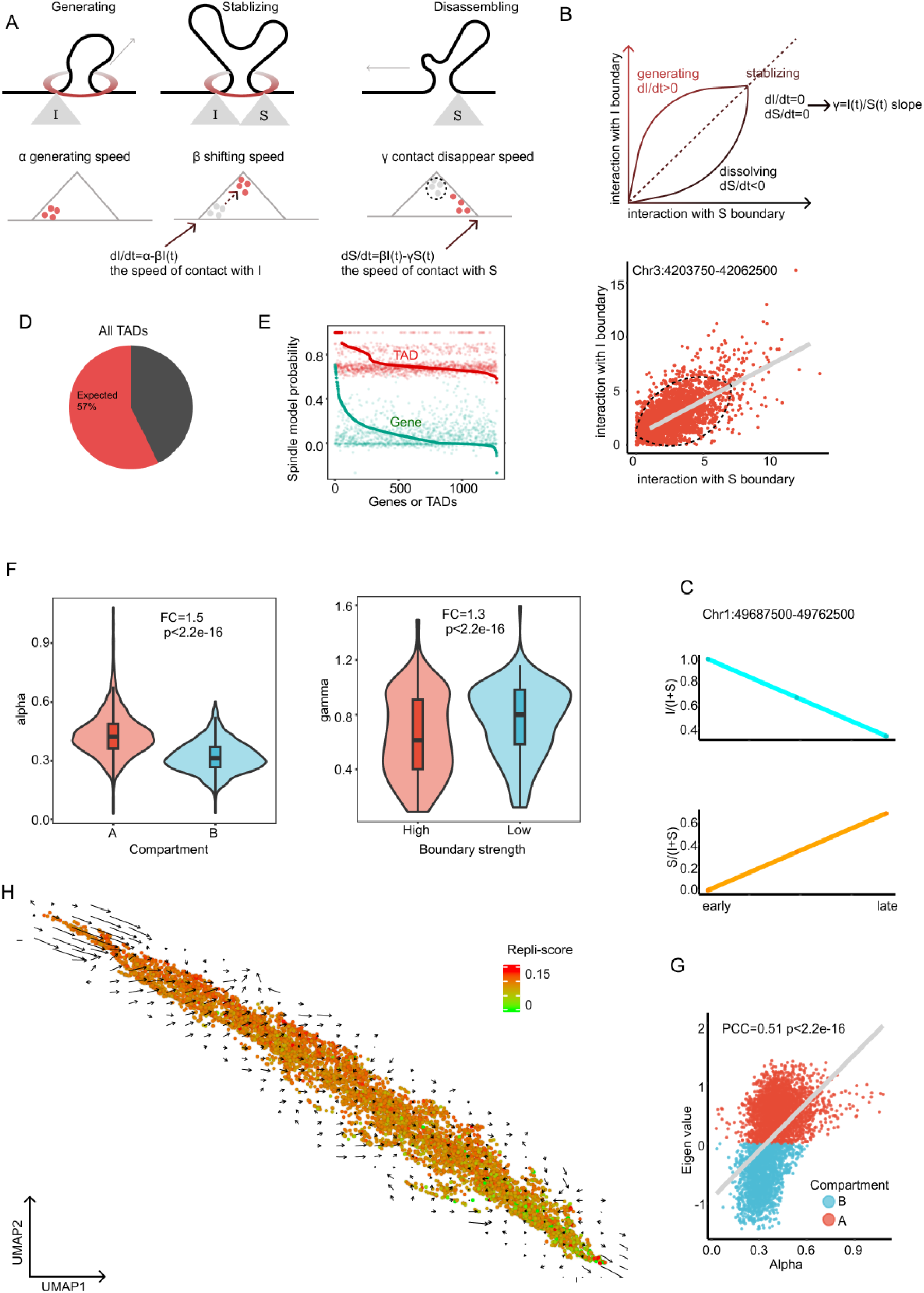
(A) This diagram illustrates our loop extrusion model, incorporating mathematical representations of these processes. (B) We’ve developed a theoretical model of the interaction between the I and S boundaries, referencing a specific region from Chr3: 4203750-42062500. (C) This graph demonstrates the average interaction ratio evolution between two boundaries along with replication time, from early to late stages. The expressions I/(I+S) and S/(I+S) represent the contact ratios of the I boundaries and the S boundary, respectively. (D) Displayed here is a pie chart indicating the percentage of the qualified spindle distribution of TADs. (E) This cumulative curve depicts the probable spindle model of TADs. (F) The violin plot showcases the alpha value in the A B compartment, with the gamma distribution in the plot signifying both high and low boundary strength. (G) This part represents the correlation of the alpha and eigen value in all TADs. (H) Lastly, the UMAP of the HEK293 cells is presented with color-coding based on the repli-score, with an arrow highlighting the loop velocity.

Once we’ve figured out the alpha gamma for each TAD, we’ll check if the velocity we’ve calculated can forecast the genome organization’s next virtual time point. We’ll then condense this information into a 2D UMAP to highlight the upcoming time point for the cell state. Within this UMAP plot, we’ll use color gradients to denote replication scores, transitioning from early to late replication stages (Figure.2H). When we took a look at the loop velocity, 60% DNA movement directions met our expectations (Sfig.4A). We then estimated the expected velocity direction and compared that with the actual direction in real cells. Our findings showed a significant correlation, with 80% of the predicted movement directions lining up with what we expected. This suggests that the alpha gamma can effectively capture and predict the state of biological processes.

In conclusion, our studies indicate that our dynamic models, integrating loop velocity, can efficiently predict cellular development direction in cultured cells. These parameters with velocity could provide a more dynamic representation for TAD.

### Perform DropHiChew for spermiogenesis cell in testis

RNA velocity has gained acceptance due to its translation activity. The key question for loop velocity is understanding its role in driving phenotypes. To examine how loop velocity impacts cell differentiation over time, we selected the mouse testis spermiogenesis model due to its rapid phenotype changes. This model also conveniently omits the mitosis process, which allows us to study loop velocity’s influence on spermiogenesis without the complication of cell cycle-related genome compaction. An added benefit of utilizing spermiogenesis cells is their unidirectional cell differentiation. They transition from diploid cells, like pachytene spermatocytes, into haploid cells such as round spermatids and elongating spermatids (Figure 3.A).

**Figure. 3.**
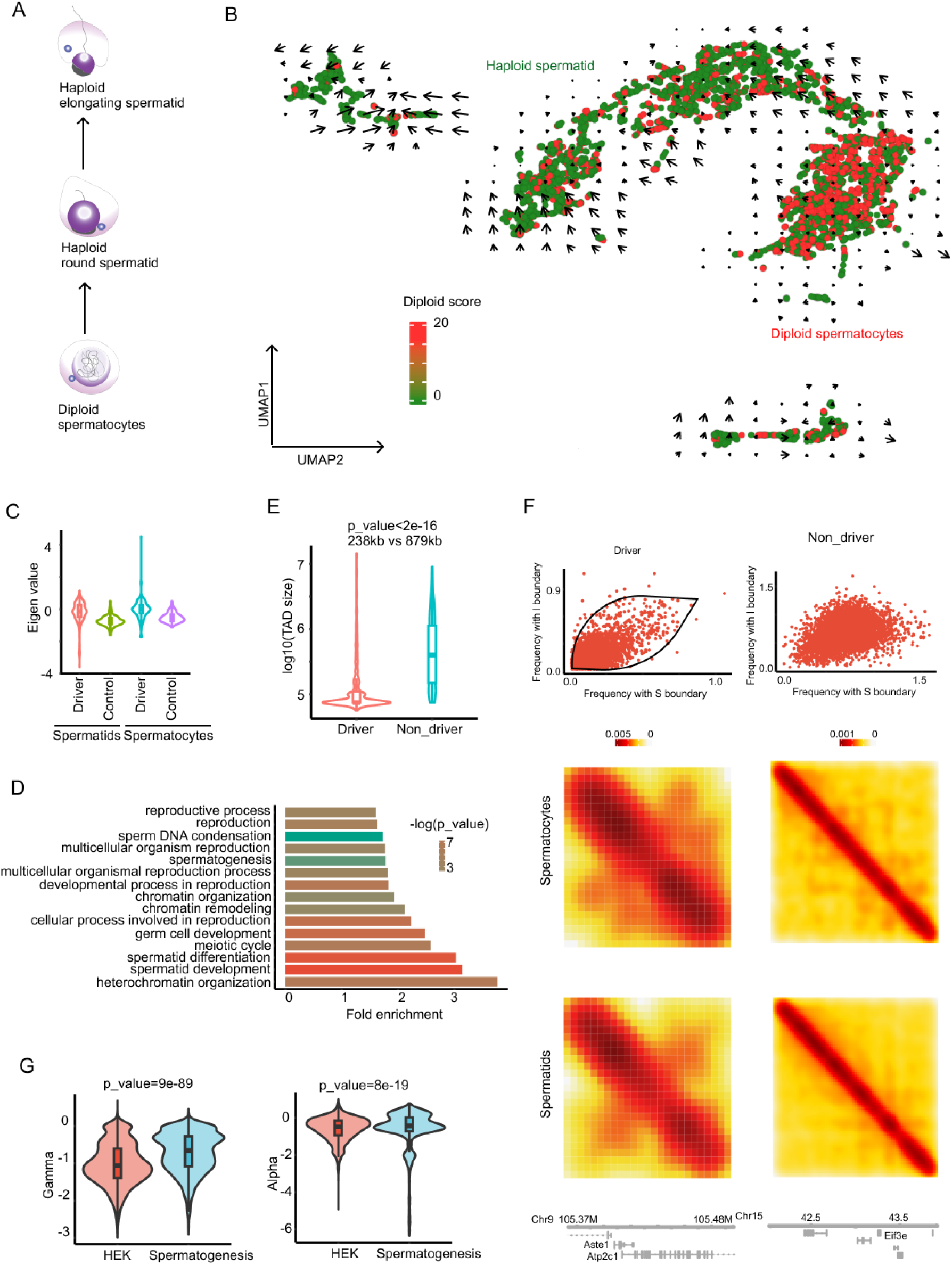
(A) Diagram showing the process of spermiogenesis. (B) UMAP of spermiogenesis cells where color denotes diploid scores. Higher scores represent pachytene spermatocytes or earlier cells, while lower scores represent haploid spermatids. Arrows are calculated using loop velocity. (C) Eigen value of the driving TADs and control non-driving TADs. (D) GO term analysis of first 1000 genes (highest likelihood) within the driving TADs. (E) Comparison of TAD size distribution between driving TADs and control non-driving TADs. (F) Examples of driving and non-driving TADs in relation to boundary contact distribution, contact maps in spermatocytes and spermatids, and gene tracks. (G) Gamma and alpha distribution of the driving TADs in HEK293 cells and spermiogenesis cells.

Following our established protocol, we prepared a single cell suspension from four adult testes and performed DropHiChew on these spermiogenesis cells. The cell filter was configured according to the method section.

Our experiment led to the detection of around 5618 cells, each with an average of 74k contacts and a duplication rate of 67%. With a valid pair ratio of approximately 35.8%, the results were comparable to our earlier assessment with HEK293T.

We then utilized the UMAP method to map and cluster cells to evaluate the effectiveness of DropHiChew in cell type classification and velocity prediction (Figure3.B). Without scRNA-seq, it becomes a challenge to use structural properties for reliable cell type classification. A key feature of meiosis is the significant change in chromosome copy number from diploid to haploid. By using the diploid scores, we can confidently distinguish between diploid spermatogenic cells (such as pachytenes and earlier cells) and haploid spermatids (round and elongating spermatids). However, we could not confidently differentiate the round spermatids and elongating spermatids.

In order to better ascertain the accuracy of the cell types, we merged the haploid spermatids and compared the contact map with control bulk samples that were FACS sorted. We observed a striking similarity in eigenvalues and insulation scores, which demonstrates the high quality of our data (Sfig3.A). Interestingly, we observed an increase in the frequency of long-distance interactions (greater than 20kb) in round spermatids(Sfig3.B), a finding that aligns with previously published results. This could be a result of genome compaction during spermiogenesis.

Once we had a clear idea of the expected cell development direction, we incorporated loop velocity into our spermiogenesis data. This allowed us to determine if we could predict the development direction accurately.

### The loop velocity accurately predict the development direction of the spermiogenesis

We’ve identified 2986 Topologically Associating Domains (TADs) within the spermiogenesis cells. Our research indicates that half of these TADs show an I_S plot spindle score above 0.4, implying a significant number of suitable objects for determining loop velocity. Our analysis suggests that loop velocity, when examined using spermatogenesis data, is a reliable predictor of the expected meiosis direction, transitioning from diploid spermatogenic cells to haploid spermatids (Figure3.B). Interestingly, prior research noted an unexpected direction when using scRNA-seq, potentially due to the short reads from scRNA underestimating the nascent RNA portion^8^. Given the strong predictability of spermiogenesis development, we performed a global quantification to assess our accuracy. The results demonstrated that loop velocity accurately predicted the expected DNA movement direction over 70% of the cases (Sfig.4A), outperforming the velocity of scRNA-seq^8^.

The process of spermiogenesis serves as an excellent model for understanding the concept of loop velocity, given the significant changes it induces in phenotype and gene expression. While RNA velocity is generally more accepted by the scientific community, this is largely because it’s understood that RNA influences cell development by translating into proteins that affect phenotype. To truly grasp the biological rationale of loop velocity in cellular prediction, it’s crucial to identify key TADs that drive development and investigate their functionality. It seems that the speed or direction of prediction arrows is influenced by the weight of each TAD’s predictive functions. Existing research implies that the expression of certain RNAs, especially those showing spindle-like dynamics (referred to as spindle model likelihood), may carry significant weight in these predictive functions^9^. Using similar methods, we’ve identified these key TADs as ‘driver TADs’.

We’ve identified approximately 1000 ‘driver TADs’, with the xxx gene located within them (likelihood>0.7) (Sfig.3C). The driver TADs exhibit a significantly higher eigenvalue than the control non-driver TADs, which suggests they may be active in transcription (Figure3.C). Interestingly, most of the driver genes are found in the sex chromosome and play a role in spermatogenesis (Sfig3.D). Upon closer examination, we found a 64% overlap with previously identified spermatogenesis driver genes (Sfig3.D). These genes show a distinct transcription pattern, contributing significantly to spermiogenesis progression. This strengthens the idea of the transcriptional activity of the driver TADs. Furthermore, it suggests that driver TADs might influence biological systems through their internal genes. To better understand their biological implications, we conducted a Gene Ontology (GO) term analysis on the genes located within the driver TADs (Figure3.D). The majority of these genes are involved in spermatogenesis-related processes, such as spermatid development and sperm DNA condensation. Interestingly, unlike housekeeping genes in cells that express steadily, these spermatogenesis-related genes vary in expression in different cells depending on the process. Overall, these findings suggest that driver TADs may enhance cellular development by promoting the expression of internal genes.

Driver TADs, if they correlate strongly with internal driver gene expression, should fluctuate in sync with gene expression, potentially driving the process of spermatogenesis. Our comparison of driver and non-driver TADs showed that driver TADs are significantly smaller, with a median size of 237kb (Figure.3E). This is anticipated, as larger TADs tend to maintain stable structures across different cell types, as indicated by previous research. On the other hand, we propose that smaller TADs may have a greater capacity for movement, which could facilitate internal gene expression.

Examining the driver TAD of two spermatogenesis stages revealed that the genome within these TADs experienced considerable movement around the boundary, which aligns with our expectations based on spindle movement (I_S plot) (Figure3.F, Sfig3E). In contrast, genomes within non-driver TADs displayed random, scattered movement. While these movements could be indirectly inferred from the alpha gamma factors, non-driver TADs couldn’t calculate the alpha gamma.

We conducted a study comparing driver TADs influencing spermiogenesis with those affecting the HEK293T cycle, to understand the volatility of motion. We found a significant difference (p_value<2e-16) with a 1.3fold change in the gamma (Figure.3G). It’s important to note that the gamma, which is the slope of the spindle model in the I_S plot, is inversely proportional to the insulation score. Therefore, an increase in gamma suggests larger and quicker shifts in genome movement. This indicates that driver TADs involved in spermatogenesis undergo more variable changes than those associated with the cell cycle. This could potentially explain why the speed arrows in the HEKs were noticeably shorter than in the spermiogenesis model, due to the smaller gamma. Therefore, driver TADs are smaller and have a higher amplitude of DNA motion, which could possibly facilitate dynamic internal gene promotion during spermiogenesis.

However, we still need direct evidence that dynamic DNA movement directly relates to gene expression, which is something we could explore in the multiomics 3C data below.

In essence, loop velocity is key in forecasting spermatogenesis progression and pinpointing the leading TADs in chromatin remodeling. It’s quite likely that these driver TADs modified internal genome structures to encourage cell differentiation by changing gene expression within the TADs.

### The loop velocity accurately predict the development direction of the embryonic stem cells

To better understand the link between chromatin movement and gene expression, it’s beneficial to have single cell data that includes both RNA and chromatin conformation. The HiRES technology, as previously described, offers embryonic data that sequences RNA and HiC in single cells^15^. This dataset is ideal for examining the relationship between RNA velocity and loop velocity. Additionally, the embryonic stem cell data, gathered over different embryonic days, provides an opportunity to verify the precision of velocity prediction by embryonic days.

We have downloaded both RNA and HiC embryonic stem cell data for embryonic days 7.0 (E7.0) to E11.5. Around 7469 cells passed filters for both data types, yielding 2.8 billion DNA contacts. The embryonic day can be used as a time indicator to evaluate the accuracy of our predictions^15^. We implemented the loop velocity on the embryonic stem cell UMAP. Some directions align correctly, transitioning from the early to the later embryonic days (indicated by the black dash square)(Figure.4A). However, the map’s central direction doesn’t appear to be accurate. Upon detailed analysis, we observed that this UMAP region amalgamates both early and late embryonic cells, which are not distinctly segregated. This factor could potentially account for the diminished performance.

**Figure. 4.**
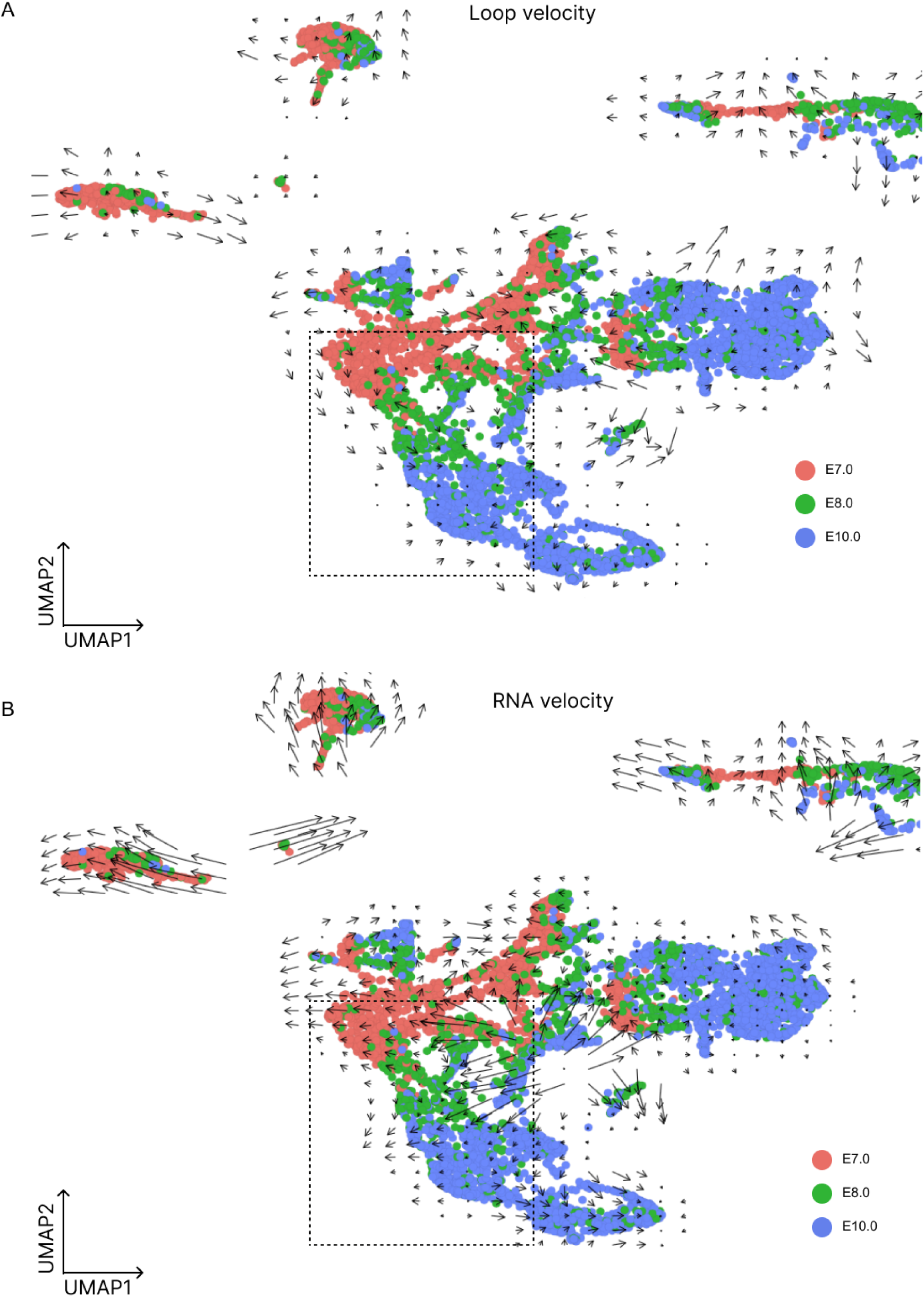
(A) Here’s a UMAP of embryonic cells from various stages. The colors represent embryonic days 7.0, 8.0, 10.0. The arrows, calculated via loop velocity, illustrate this. (B) These arrows were determined using RNA velocity.

We’ve also run the RNA velocity test for comparison (see Figure.4B). The findings suggest that RNA velocity didn’t perform as well as loop velocity, even in areas with clear separation (indicated by the black dashed square). When we calculated the global direction in loop velocity, we found that more than half of the DNA movement predictions were heading in the same direction in the anticipated cells (Sfig.4A).

Our team performed a GO term analysis within the driver TADs to pinpoint the specific TADs impacting prediction speed (Sfig.4B). We observed a heightened enrichment of genes linked with the immune system and monocyte differentiation. These findings imply that the dynamic models are capable of accurately forecasting the cell differentiation process.

Considering that driving TADs may encourage cell state via internal gene expression, we still lack direct evidence. Embryonic studies, using simultaneous RNA and 3C data from the same cell, have provided clearer insight into the potential relationship between DNA movement in TADs and gene expression. We’ve pinpointed loop extrusion from the I boundary to the S boundary. It seems that gene expression amplifies when interactions mainly occur at the I boundary (upper curve). Conversely, when chromatin is primarily at the S boundary (lower curve), gene expression diminishes (Sfig.4C). We notice significant expression changes when the chromatin shifts between these two boundaries (Sfig.4D). Further analysis shows that this pattern is consistent in about 92% of TADs, suggesting a strong correlation between gene expression and chromatin movement.

In summary, we’ve introduced the concept of loop velocity through the simple experimental system of DropHiChew. This novel approach allows us to introduce the dynamic parameters alpha gamma to describe the dynamic movement of the TAD and predict the direction of cellular development. Our loop velocity model has shown high predictive accuracy across various cell types, including HEK293 cells, spermatogenesis cells, and embryonic stem cells. Intriguingly, our findings suggest that loop velocity not only predicts cellular development direction, but also identifies driver TADs that influence this development through changing the gene expression within TADs. This research provides valuable insights into the dynamics of chromatin architecture within individual cells, offering a valuable tool for future studies in cellular development.

## Discussion

In the results section, we present the DropHiChew - the first commercial platform to incorporate single cell 3C experiments. Our unique approach to post-PCR enrichment technology significantly improves data usage, achieving 4x the performance of Dip-C and making it more accessible to general labs. Simultaneously, we introduce the concept of loop velocity, an innovative method to predict cellular development direction by examining the dynamics of Topologically Associating Domains (TADs). With its accuracy in predicting cellular development direction, loop velocity also helped us identify key TADs driving development, providing fresh insights into the dynamics of chromatin architecture within individual cells.

We need to address a critical shortfall in our 3C experiment - the effectiveness of Tn5 in crosslinked cells. Our observations showed that Tn5 underperforms in this context, leading to a loss of many contacts. Furthermore, the natural process of Tn5 may also result in losing half the contacts due to homologous attachment. In response to this, other researchers have refined the problem and developed a method for the imputation of HiC contacts, allowing for ample single-cell contacts for most uses.

Moreover, given the use of Tn5, it’s natural to question whether Tn5 might create a bias towards accessible regions. However, after comparing the HiChew(open/close ratio 0.1), a ligation-based method, and DropHiChew (open/close ratio 0.02), a Tn5-based method, we didn’t observe the contact bias towards accessible regions. In fact, we found that Tn5 exhibits less preference for the accessible region in the DropHiChew. This is likely due to heated SDS being able to resolve most compact chromatins. The similarity of the contact maps further supports this point of absence of bias towards accessible chromatin. However, there are differences that exist. We believe that the limitations of Tn5 should be given more consideration in future 3C research.

The other shortage in this experiments, we used the DpnII-Dam methyaltransferase to perform the DropHiChew. Therefore, the digestion enzyme selection need the corresponding the mehtylattransferase to labeling the ligation scar in order to perform the post PCR enrichment. These combination selection is around 20 different enyzmes, which might limited your choice of enzyme. Moreover, multiple enyzme digestion is not available for this method. therefore the high resolution maps is not achievable in the DropHiChew.

In this project, we recognize that a reduced contact density could limit our ability to observe detailed chromatin changes. Deep sequencing, while valuable for observing intricate structures, may hinder many labs from processing numerous samples due to cost. So, the challenge we’re addressing is how to best utilize shallow sequencing of single cell HiC and take full advantage of this accessible platform. To address this, we created a new algorithm, “loop velocity”, that remains reliable even with fewer contacts and doesn’t necessitate imputation. Furthermore, with careful application, the alpha and gamma could potentially infer the compartment eigenvalue and insulation score. Most notably, the movement momentum now carries biological significance in the development analysis.

In the context of loop velocity applications, we discovered that the key to accurate results lies in UMAP clustering. However, both our research and previous publications have shown that 3C data doesn’t differentiate as well as RNA-seq data at the single-cell level^15,16,20^. If 3C data can’t discern subtle changes in UMAP, it could lead to errors in loop velocity. Consequently, there’s a pressing need for more sophisticated clustering methods specifically designed for 3C. Fortunately, recent advancements in the field have seen the development of simultaneous methods for detecting RNA expression and 3C in single cells. Alternatively, integrating two different datasets can also provide better resolution in cell clustering, which forms a solid foundation for loop velocity analysis. Moreover due to our system is deeply customzied to 10X system, therefore this method could easily adapt to the 10X Chromium Single Cell Multiome ATAC + Gene Expression system, which could simoutaniously detect the 3C and RNA expression in the single cell.

The interplay between genome folding in cell cycles and cell differentiation can be intricate, yet it’s essential, particularly in embryonic cells where these processes are happening at the same time. We’re primarily exploring whether cell differentiation or the cell cycle is more influential in our predictions. As per our driver TAD analysis, TADs associated with cell differentiation appear to hold the most predictive value. This could be attributed to these TADs typically having higher gamma factors than those linked to the cell cycle. So, in our research, it seems that TADs related to cell differentiation are the key determinants of prediction direction. Additionally, we predict local speed using local neighboring data points on the UMAP. If the two kinds of driver TADs don’t vary simultaneously, they shouldn’t interfere with each other in the analysis.

In the loop velocity theory, we discussed single-sided extrusion models. Yet, some scientists might suggest that TAD forms from both sides^21^. To address this, we’ve also developed a two-sided extrusion model, which you’ll find in the supplemental notes for further exploration. We used statistical tests to examine the contact density on both sides to determine if these TADs resulted from one or two-sided extrusion. Interestingly, we found that 70% of TADs rely on one-sided extrusion, with the remainder depending on two-sided extrusion. The one-sided model could cover most of the TADs in our downstream analysis.

In previous studies involving the RNA velocity field, scientists identified noise issues resulting from single cell RNA sequencing that impacted the accurate determination of nascent RNA. As a result, a variety of machine learning algorithms were developed to counterbalance this noise and improve prediction accuracy. In our loop velocity, we experimented with the Expectation-Maximization (EM) algorithm to evaluate potential performance enhancements. Our results, however, did not indicate improved performance. In earlier RNA velocity research, it was identified that preset parameters derived from a perfect spindle model (nascent/degrading RNAs) can render the EM algorithm unnecessary. Our previous research utilized long-range sequencing for precise nascent RNA identification, leading us to discover that sample quantile regression proved to be robust under various conditions. Our findings from the loop velocity study indicated that changes in the I-S boundary contact formed an ideal spindle model. Therefore, simple quantile regression demonstrated sufficient robustness, eliminating the need for more complex machine learning algorithms.

In general, the loop velocity and the experimental DropHiChew provide a manageable approach for most scientists to delve into DNA dynamics.

## Method

### Cell Culture

We cultured both HEK293T, a human embryonic kidney cell line expressing a mutant SV40 large T antigen, and 3T3, a line of mouse embryonic fibroblasts, in DMEM (Gibco, no. 11965092). This was supplemented with 10% FBS (Gibco, no. 10099141) and 1× penicillin-streptomycin (Gibco no. 15140122), and maintained at 37 °C under 5% CO2.

### Fixation protocol

The cells we collected were rinsed twice with cold PBS. We then cross-linked them in PBS containing 1% FA (Thermo, No. 28908) for 10 minutes at room temperature. Afterwards, we neutralized the reaction with 250 mM glycine for 5 minutes at room temperature. Finally, we rinsed the cells twice again with cold PBS.

### Cell membrane penetration and chromatin decondensation

We resuspended the cell pellet in 500 μl of lysis buffer (containing 50 mM HEPES pH 7.4, 1 mM EDTA pH 8.0, 1 mM EGTA pH 8.0, 140 mM NaCl, 0.25% Triton X-100, 0.5% IGEPAL CA-630, 10% glycerol, and a 1× protease inhibitor cocktail). We then let this sit on ice for 10 minutes before centrifuging it at 500g for 3 minutes at 4°C. The sample was then washed once with wash buffer (10 mM Tris-HCl pH 8, 1.5 mM EDTA, 1.5 mM EGTA, 200mM NaCl, 1× protease inhibitor cocktail) and centrifuged again at 500g for 3 minutes at 4°C. We removed and discarded the supernatant, and gave the pellet one more wash with 1× DpnII buffer.

The supernatant was carefully removed once more, and the pellet was then resuspended in 199 μl of 1.2×DpnII buffer, inclusive of 0.1% SDS. Following a thorough stirring, the pellet was treated in a 65°C metal bath for a duration of 10 minutes, swiftly followed by a 2-minute cooling period on ice. The process concluded with the addition of 10 μl of 20% Triton X-100, accompanied by a 15-minute incubation at 37°C with consistent agitation.

### Enzyme Digestion and ligation

We started by adding 200U DpnII (NEB, R0543L, final concentration 1U/μl) to the previous reaction mix, giving it a good stir, and letting it react overnight at 37°C. Then, we added 2 μl of the digestion reaction mix to 7 μl NF water and 1 μl proteinase K, and let it all react at 65°C for an hour to break down the crosslinks. We checked the size of the DNA fragments using 1% gel electrophoresis.

Next, we deactivated DpnII by heating it at 65°C for 20 minutes. We then got rid of the supernatant through centrifugation and resuspended the pellet in 195 μl 1×T4 DNA ligase buffer.

We added 0.5μl of 2000 U/μl T4 DNA ligase (ABclonal, RK21500) to achieve a final concentration of 10 U/μl. After a thorough mix, we let it react overnight at 16°C.

Finally, we took another 2 μl of this solution, added it to 7 μl NF water and 1 μl proteinase K, broke down the crosslinks at 65°C for 1 hour, and confirmed the size of the bands using 1% gel electrophoresis.

### ChromiumNext GEM Single Cell ATAC

Nuclei were passed through a 40 µm (pluriSelect, no. 43-10040-40) and a 10-μm (pluriSelect, no. 43-10020-60) mesh filter. Afterwards, the instructions from the 10x Single Cell ATAC Operation Manual were adhered to.

### Methylation labeling and immunoprecipitation

The outcome of the previous procedure was amplified via PCR using new P5 and P7 primers to substitute the GTAC site and index at the P5 end with CATC or others (P5 5’ - /phos/ AATGATACGGCGACCACCGACATCTACAC - 3’ P7 primers with phosphorothioate bond 5’ - C*A*A*G*C*AGAAGACGGCATACGAGAT - 3’). Next, we added 1 μg of DNA to the reaction, including 1 μl Lambda Exonuclease, 5 μl 10× Lambda Exonuclease Reaction Buffer, and water up to 50 μl. This mixture was then incubated at 37°C for 30 minutes.

Afterwards, we purified the product and turned the remaining single strand into a double-strand using a short RN1 primer (RN1 primer 5’-TCGTCGGCAGCGTCAG-3’). We then added 500 ng of this purified, extended DNA to a reaction system composed of 5 μl 10×dam Methyltransferase Reaction Buffer, 0.25 μl 32 mM SAM, 1 μl dam methyltransferase, and enough water to make up 50 μl in total. We incubated this at 37°C for 1 hour to mark the GATC site m6A methylation.

Next, we treated the Protein A/G beads: we rinsed 10 μl Protein A/G beads twice with 1X PBST buffer containing 0.1% Tween 20, then re-suspended them in 50 μl SuperBlock™ Blocking Buffer and incubated at room temperature for 15 minutes. We then washed the PA/G beads twice with 1×IP buffer and re-suspended them in 48 μl 1×IP buffer. We added 2 μl m6A antibody to this and allowed it to rotate at 4°C overnight.

Finally, we washed the Protein A/G beads pre-bound with antibody with 1×IP buffer and re-suspended them in 50 μl 1×IP buffer for later use. We diluted the purified DAM-labeled DNA to 40 μl with EB buffer, denatured it at 95°C for 5 minutes, and immediately placed it on ice for 2 minutes. We then added 10 μl of pre-chilled 5× IP buffer and 50 μl of pre-bound anti-m6A Protein A/G beads to the deformed DNA and mixed thoroughly. This was incubated at 4°C with rotation for 2 h.

We started by placing the beads on a magnetic stand and removing the supernatant. Next, we gave the beads a quick wash with pre-chilled medium stringency RIPA buffer. The beads then received two washes on ice with pre-chilled high stringency RIPA buffer. After pre-chilling, we washed the beads once more with stringent RIPA buffer and gave them two final rinses with cold 1× IP buffer.

We then resuspended the magnetic beads in 20 μl EB and added Qiagen protease to achieve a final concentration of 0.05 U/ml. The mixture was incubated at 50°C for 30 minutes and then at 70°C for 15 minutes.

Following this, we added 30 μl of a PCR mix (25 μl KAPA HiFi HotStart ReadyMix, 2 μl P5 primer, 2 μl P7 primer, 1 μl water) and ran 10 cycles of amplification on a PCR instrument. Finally, we purified the library with 1× Ampure XP beads.

### Library Circulation and Sequencing

The library was prepared using the MGIEasy Cycling Kit (MGI, No. 1000005259), as per the kit instructions. The product was then processed with the MGISEQ-2000RS High-throughput Sequencing Kit (PE100) (MGI, No. 1000012554), for the creation of DNA nanoballs. These were sequenced using the PE100+100+10+16 mode on the MGISEQ-2000 platform and T7 MGI platform.

### Preprocessing of DropHiChew datasets

We began by sorting the raw reads for each cell with our in-house script. Then, we aligned single-cell Hi-C paired-end reads to either the hg19 or mm10 reference genome using HiC-Pro (v.3.1.0, default settings). This gave us HiC-Pro filtering and alignment metrics, valid pairs, and contact matrices. We corrected the matrices with ice_norm. We created contact matrices for each chromosome at resolutions from 10kb to 1Mb for more detailed analysis.

To estimate the number of valid cells in the DropHiChew data, we used a barcode rank plot to pinpoint the steep drop-off pattern that distinguishes valid cells from background noise. We plotted all detected DropHiChew barcodes in descending order by the number of nonduplicated valid pairs linked to each barcode. The R package kneedle helped us identify the transition between valid cells and non-cell barcodes. We then used the data associated with the valid cells for additional analysis.

### Comparison of pseudo bulk chromatin architecture datasets

We’ve taken a look at important metrics such as the cis/trans ratio, valid pair ratio, valid pair dup, among others from the HiC-Pro statistical outputs. To give you a clearer picture, the valid pair ratio is calculated by dividing the valid pairs (before duplication removal) by the reported pair from HiC-Pro. This method effectively illustrates the enrichment efficiency. On the other hand, the valid pair dup refers to the duplicated portion of valid pairs, as identified by HiC-Pro, providing an indication of the level of sequencing saturation.

To study the pattern of the contact matrix, we initially compiled the nonduplicated valid pairs from every cell and transformed them into mcool files with the help of the cooler tool (v0.8.2). Following this, we utilized the HiCExperiment (v1.2.0) R package to conduct a comparative analysis in pairs at various contact map resolutions. This analysis provides a deeper understanding of Hi-C features at the levels of the entire genome, specific chromosomes, topological associated domains (TAD), and chromatin loops.

We evaluated the correlation of contact matrices at both chromosomal and TAD levels by utilizing genome-wide eigenvector scores (compartment score) and TAD insulation score, as determined by Cooltools (v0.4.1). Following this, the correlation was calculated using our own R scripts and the Pearson methods. To conclude, the contact distance decay curve was determined using HiCExplorer (v3.7.2).

### DropHiChew collision rate estimation

The data from the snHiChew HEK293T-NIH3T3 mix was processed using a combined genome of hg19 and mm10. This was done using the standard snHiChew preprocessing methods. We utilised nonduplicated valid pairs in the cell rank plot to spot a sharp decline, identifying the empty cell barcodes. If a minimum of 89% of nonduplicated valid pairs were linked to hg19, we tagged the valid cell barcodes as HEK293T. We did the same for NIH3T3, tagging them if 89% of pairs were linked to mm10. Any remaining valid cell barcodes were considered as doublets.

### Clustering of DropHiChew data in HEK293T-NIH3T3 mixture data

We’ve grouped the corresponding pairs related to the annotated HEK293T and NIH3T3 cell barcodes. The dimensionality reduction calculations were carried out using Higashi ^22^, with standard settings in place. For UMAP clustering, we applied the first 15 principal components of the Higashi embeddings. The parameters were set to “n_neighbors=5, min_dist=0.01, metric=’correlation’”.

### Comparison of single cell chromatin architecture datasets

We’ve taken a look at how DropHiChew’s chromatin architecture data stacks up against data from snHiChew, Dip-C and snHi-C. These are all well-respected single-cell chromatin conformation capture technologies. The datasets we used for this comparison come from the National Center for Biotechnology Information (NCBI) Gene Expression Omnibus (GEO), under the codes GSE94489 and GSE146397.

We utilized identical preprocessing and HiC-Pro metrics-based benchmark procedures for Dip-C, snHi-C, and snHiChew as we previously delineated. In order to evaluate sequencing yield efficiency, we reduced raw reads to diverse levels: 25k, 50k, 100k, 200k, 400k, 800k, 1.6M, 3.2M, 6.4M, and 12.8M. We strictly included cells whose read numbers surpassed these set thresholds.

Afterwards, we figured out the count of unique valid pairs per read for each threshold. We did this by fitting the data points to a saturation curve, using the model Y = B max * X/(Kd + X). To wrap things up, we gauged the read number for sequencing saturation for each single-cell chromatin conformation capture technology.

We examined the false positive rate by observing unexpected interactions between mitochondrial DNA and nuclear DNA. The false positive rates for each distinctive valid pair identified by HiC-Pro in every valid cell of the snHiChew and Dip-C datasets were calculated, as per the method outlined in Trac-looping ^23^.

### In silico HEK293 cell phasing over the cell cycle

We performed cell-cycle analysis using the method outlined in a previous study ^24^. Essentially, we utilized the HEK293 2-phase Repli-seq dataset (4DNESSV33VOL, 4DNESH4XLJCW) sourced from the 4D Nucleome Data Portal to label the early/late repli-score ratio for each cell. A higher early/late repli-score ratio signifies a cell closer to the early S-phase of the cell cycle.

### In silico testis cell phasing over the sex chromosome ratio

We performed an analysis of testis cell phasing, using the ratio of sex chromosome to autosome. Essentially, we labelled the sex chromosome ratio for each cell by dividing the number of fragments on chromosome X by the sum of chromosome X and chromosome 7 fragments. A sex chromosome ratio closer to zero signifies that the cell is more likely in the round or elongating spermatid phase of spermatogenesis.

### DropHiChew velocity input feature extraction

We took the pooled single-cell valid pairs, converted them into mcool files with a resolution of 25kb, and balanced them using cooler (v0.8.2). To identify pseudo bulk (pooled single cell) TAD boundaries, we utilized hicFindTADs from HiCExplorer (v3.7.2). We set parameters to “--minDepth 100000 --maxDepth 750000 --step 50000 --thresholdComparisons 0.05 --delta 0.01 --correctForMultipleTesting fdr -p 20”.

Next, we quantified the 1D count of the inner 50k region for the left and right boundary of each pseudo bulk TAD using bedtools intersect for each cell. We then normalized the 1D count for each region in accordance with the sequencing depth.

### Statistics

We carried out a two-sided t-test on the majority of the parametric data, adhering to a normal (or log-normal) distribution. In addition, we conducted a Pearson correlation analysis on this data. For non-parametric data or data that didn’t follow a normal distribution, we utilized the Wilcoxon rank test.

### Loop velocity

Refer to the supplemental note.

## Data availability

You can find the data for this study at the following locations: <u>NCBI</u> and NCBI BioProject PRJNA1127783. We also used other public datasets from NCBI GEO, which you can access using these identifiers: ChIP-seq (CTCF ENCSR135CRI H3K4me3 ENCSR000DTU; H3K27ac ENCSR000FCH), snHi-C (GSE94489), Dip-C (GSE146397), HEK293T in situ Hi-C (GSE143465), SCA-seq (PRJNA917827). The source data come with this paper.

## Code availability

Custom scripts used in this study are available from https://github.com/genometube/DropHiChew

## Author contributions

CT designed and oversaw the experiments. CT conducted the laboratory experiments; CZ and YMX conducted the bioinformatics data analysis. All authors collectively performed the data analysis. All authors have read and approved the final draft of the manuscript.

---

**Supplemental Figure 1.**
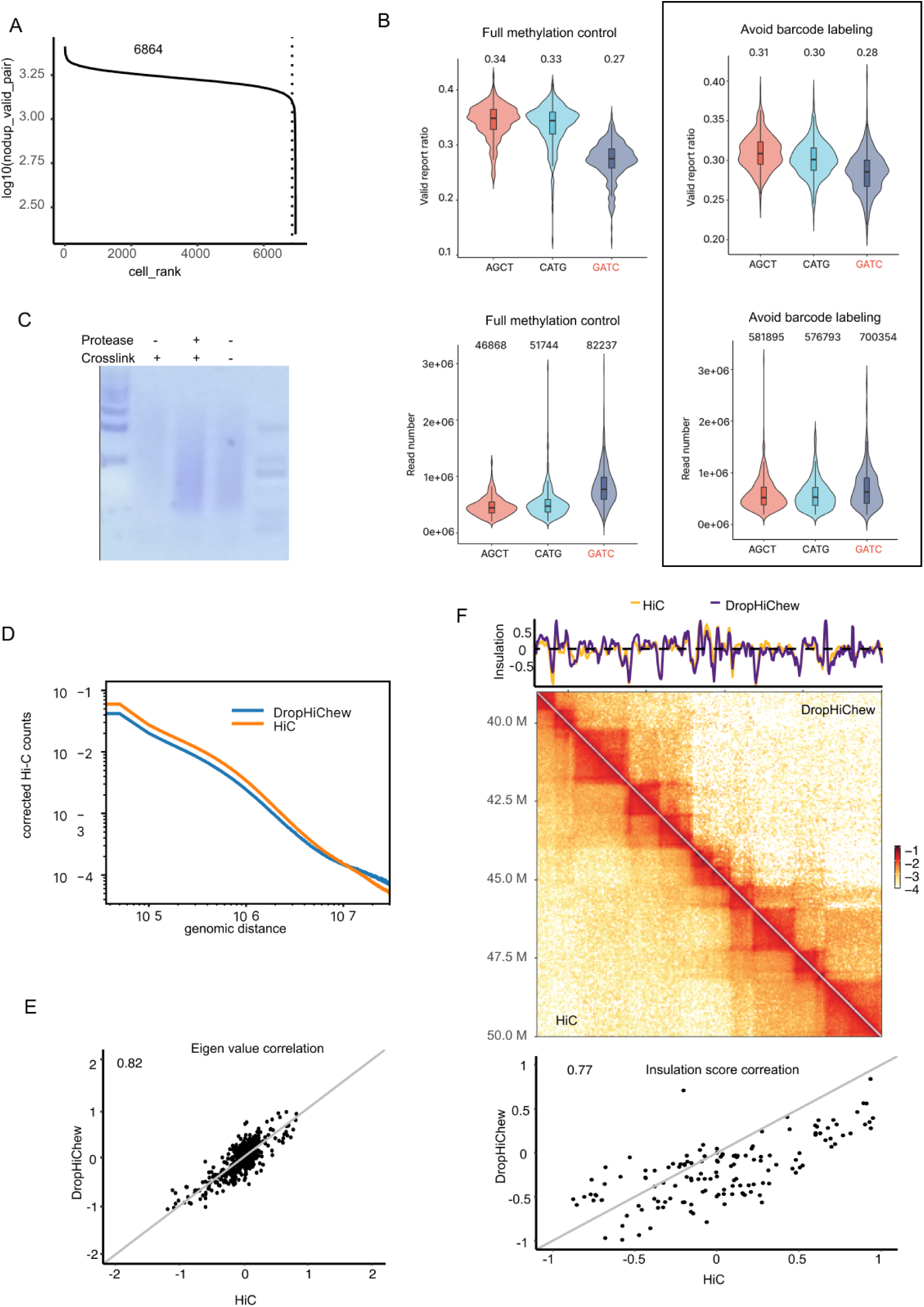
(A) Depicts the cell rank curve with deduplicated valid contacts. (B) Compares the read number and valid report ratio (valid pairs/valid reads) in the control full methylation sample and the DropHiChew, which avoids barcode labeling. (C) Illustrates the Tn5 fragmentation test on crosslinked cellular DNAs, decrosslinked cellular DNAs, and control cellular DNAs. (D) Presents the contact distance decay curve of the DropHChew. (E) Showcases the Eigen value correlation between DropHChew and HiC. (F) Compares the contact maps, insulation score fluctuation and insulation score correlation between HiC and DropHChew.

**Supplemental Figure 2.**
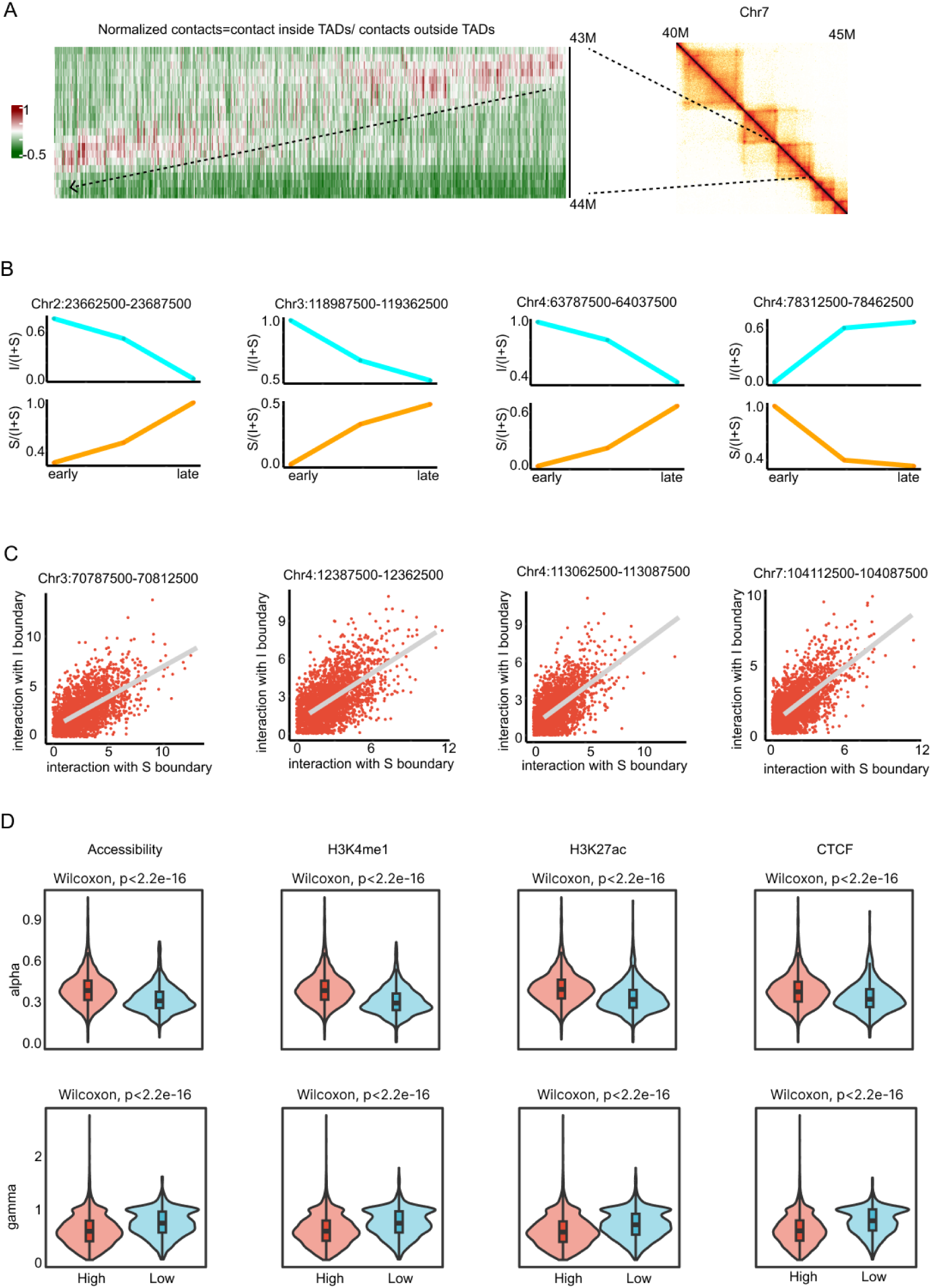
(A) This illustrates the normalized contacts plot to the 1D genome of Chr7:43M-44M. (B) It presents the boundary contact ratio of the I boundary and S boundary. (C) This is a depiction of the interaction distribution between the I boundary and S boundary. (D) Finally, it shows the alpha and gamma distribution between high and low chromatin accessibility, H3K4me1, H3K27ac, and CTCF.

**Supplemental Figure. 3:**
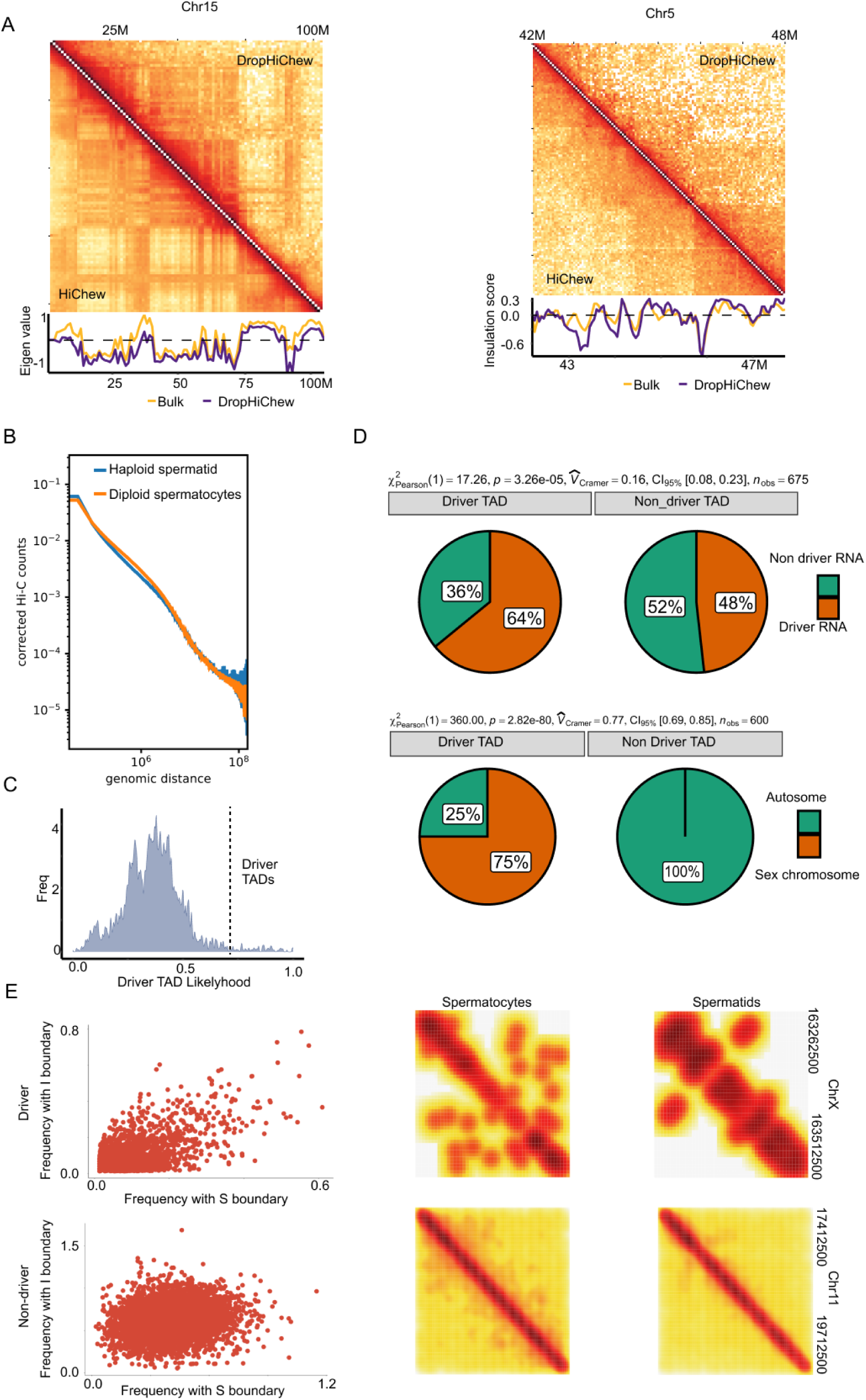
(A) Here, we compare the eigenvalue fluctuation and insulation score fluctuation of HiC and DropHiChew, as shown in the contact maps of Chr15:0-100Mb and Chr15:42-48Mb. (B) We present the distance decay curve of contact between Haploid spermatid and diploid spermatocytes. (C) We illustrate the spindle shape likelihood distribution of TADs, with likelihoods >0.7 defined as driver TADs. (D) We display the Chi-square test distribution of overlap between driver/non-driver TADs and driver/non-driver genes. (E) We provide contact maps of example regions of driver and non-driver TADs. For example, ChrX:163262500-163512500 serves as a driver TAD, and Chr11 17412500-19712500 as a non-driver TAD. Additionally, we depict the contact frequency distribution between I boundary and S boundary in all cells as point plots.

**Supplemental Figure. 4.**
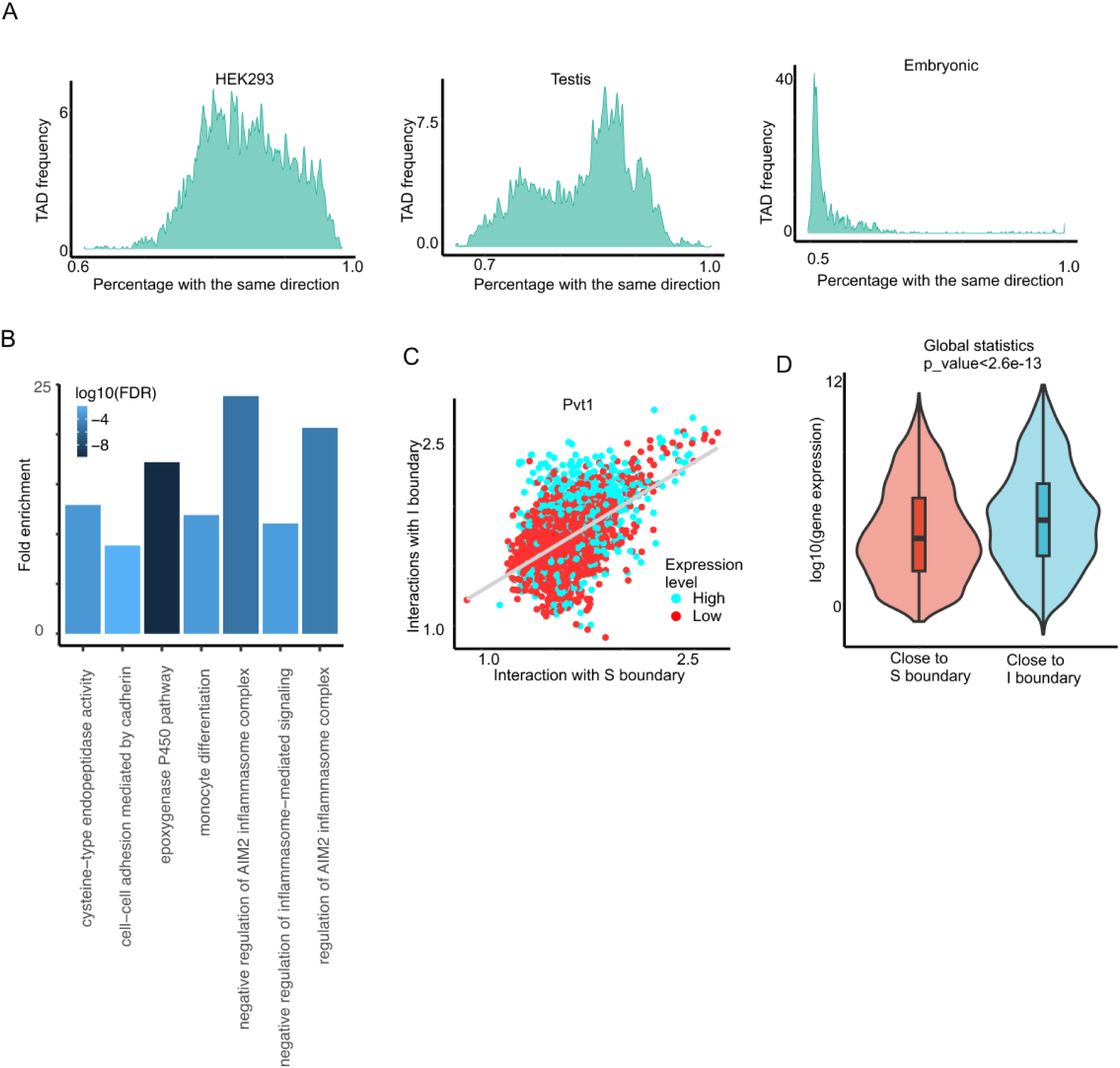
(A) We’ve compiled the TAD frequency data to forecast the correct directions of movement for the next time point in HEK293 samples, spermiogenesis samples, and embryonic samples. (B) We’ve conducted a GO term analysis of the first 1000 genes in Driver TADs. (C) We observed changes in Pvt1 abundance correlating with the DNA movement between I boundary and S boundary. (D) We have gathered global statistics of gene expression when moving closer to the I and S boundaries. (E) We’ve calculated the percentage of genes that match the pattern of gene expression, exhibiting high expression around the I boundary and lower expression around the S boundary.

